# Learning improves conscious access at the bottom, but not the top: Reverse hierarchical effects in perceptual learning and metacognition

**DOI:** 10.1101/073130

**Authors:** Benjamin Chen, Matthew Mundy, Naotsugu Tsuchiya

## Abstract

Experience with visual stimuli can improve their perceptual performance, a phenomenon termed visual perceptual learning (VPL), but how does VPL shape our conscious experience of learned stimuli? VPL has been found to improve measures of metacognition, suggesting increased conscious stimulus accessibility. Such studies however, have largely failed to control objective task accuracy, which typically correlates with metacognition. Here, using a staircase method to control this confound, we investigated whether VPL improves the metacognitive accuracy of perceptual judgements. Across three consecutive days, subjects learned to discriminate faces based on either their identity or contrast. Holding objective accuracy constant, perceptual thresholds improved in both tasks, while metacognitive accuracy diverged, with face contrast VPL improving metacognition, and face identity VPL failing to. Our findings can be interpreted in a reverse hierarchy theory-like model of VPL, which counterintuitively predicts that the VPL of low- but not high-level stimulus properties should improve conscious stimulus accessibility.

---

The relationship between subjective, conscious experience and learning remains a central topic in cognitive neuroscience (Bayne, Cleeremans, & Wilken, 2009). Learning effects however, have been largely investigated in the context of objective task performance. For instance, visual perceptual learning (VPL) describes the mechanisms whereby experience with visual stimuli improves their subsequent perceptual performance (e.g., discrimination accuracy; Fahle & Poggio, 2002). While VPL has been closely associated with non-conscious processes (Seitz, Kim, & Watanabe, 2009; Squire & Wixted, 2011; Watanabe, Nanez, & Sasaki, 2001), the exact relationship between VPL and consciousness is poorly understood. Investigating this issue, two parallel bodies of research have recently emerged. One investigates whether conscious stimulus awareness is necessary for VPL (e.g., Seitz et al., 2009; Watanabe et al., 2001), while the other investigates whether conscious stimulus accessibility can be improved by VPL (e.g., Schwiedrzik, Singer, & Melloni, 2011). Addressing the latter, we investigated whether VPL could improve the metacognitive accuracy of perceptual judgements.

Providing an account of VPL and visual consciousness is reverse hierarchy theory (RHT; Ahissar & Hochstein, 2004; Hochstein & Ahissar, 2002). According to RHT, bottom-up processing is sufficient for VPL occurring at the apex of the visual hierarchy (e.g., VPL of high-level stimulus properties), while top-down processing is needed to guide VPL to lower visual regions when greater spatial resolution is needed (e.g., VPL of low-level stimulus properties). Likewise, bottom-up processing enables conscious awareness of a stimulus’ general, global information (i.e., its gist), while top-down guided processing towards lower visual regions are needed to consciously access and become aware of its specific, local elements. Although independent lines of research support RHT in VPL (Ahissar, Nahum, Nelken, & Hochstein, 2009; see however, Watanabe et al., 2001) and visual consciousness (Campana & Tallon-Baudry, 2013), precisely how conscious stimulus accessibility interacts with the level of VPL in the presence and absence of top-down processing remains unknown. One possibility is that conscious accessibility also operates in reverse of the visual hierarchy, beginning by accessing high-level stimulus gist via bottom-up processing, and then accessing its lower-level properties via top-down processing. Thus, in the absence of top-down guided VPL, the gist of the learned stimulus can be accessed (e.g., general category), but the learned stimulus components supporting VPL cannot (e.g., precise spatial relationship between stimulus elements). Consequently, conscious accessibility fails to improve with VPL in these circumstances. Conversely, in the presence of top-down guided VPL, these learned stimulus components can be accessed, leading to improved conscious accessibility with VPL. Consistent with the latter prediction, studies investigating the VPL of simple object properties (e.g., shape) have reported accompanying improvements in measures of metacognition (Bertels, Franco, & Destrebecqz, 2012; Schlagbauer, Muller, Zehetleitner, & Geyer, 2012; Schwiedrzik et al., 2011). However, does the VPL of high-level stimulus properties fail to improve accompanying metacognition?

In the present study, we aimed to investigate whether or not the VPL of face identity, a high-level face property, fails to improve the metacognitive accuracy of perceptual judgements. Measures of metacognition however, typically correlate with objective task performance (Galvin, Podd, Drga, & Whitmore, 2003; Sandberg, Timmermans, Overgaard, & Cleeremans, 2010), a confound that has been largely overlooked in VPL studies of metacognition. Thus, any improvements in metacognition could be an artefact of uncontrolled task performance, rather than VPL per se. To control this confound, we employed a staircase paradigm during training. Using staircase methods, VPL is indexed as a decrease in the stimulus intensity (threshold) needed to maintain a given level of objective task performance (Gold, Law, Connolly, & Bennur, 2010). Such changes suggest a horizontal shift of the psychometric function corresponding to task accuracy as a function of stimulus intensity (Figure 1A). Like objective task accuracy, however, metacognitive accuracy is also thought to increase in a sigmoidal fashion with stimulus intensity (Koch & Preuschoff, 2007; Sandberg, Bibby, Timmermans, Cleeremans, & Overgaard, 2011). Assuming that metacognitive accuracy is also characterized as a psychometric function, two competing hypotheses are proposed depending on whether or not VPL improves metacognitive accuracy. If VPL improves metacognitive accuracy, this function should shift horizontally after training such that, like objective accuracy, a given level of metacognitive accuracy is maintained before and after training (Figure 1B). If VPL does not improve metacognitive accuracy, this function should remain static, resulting in metacognitive accuracy decreasing alongside stimulus threshold (Figure 1C).

**Fig. 1.**
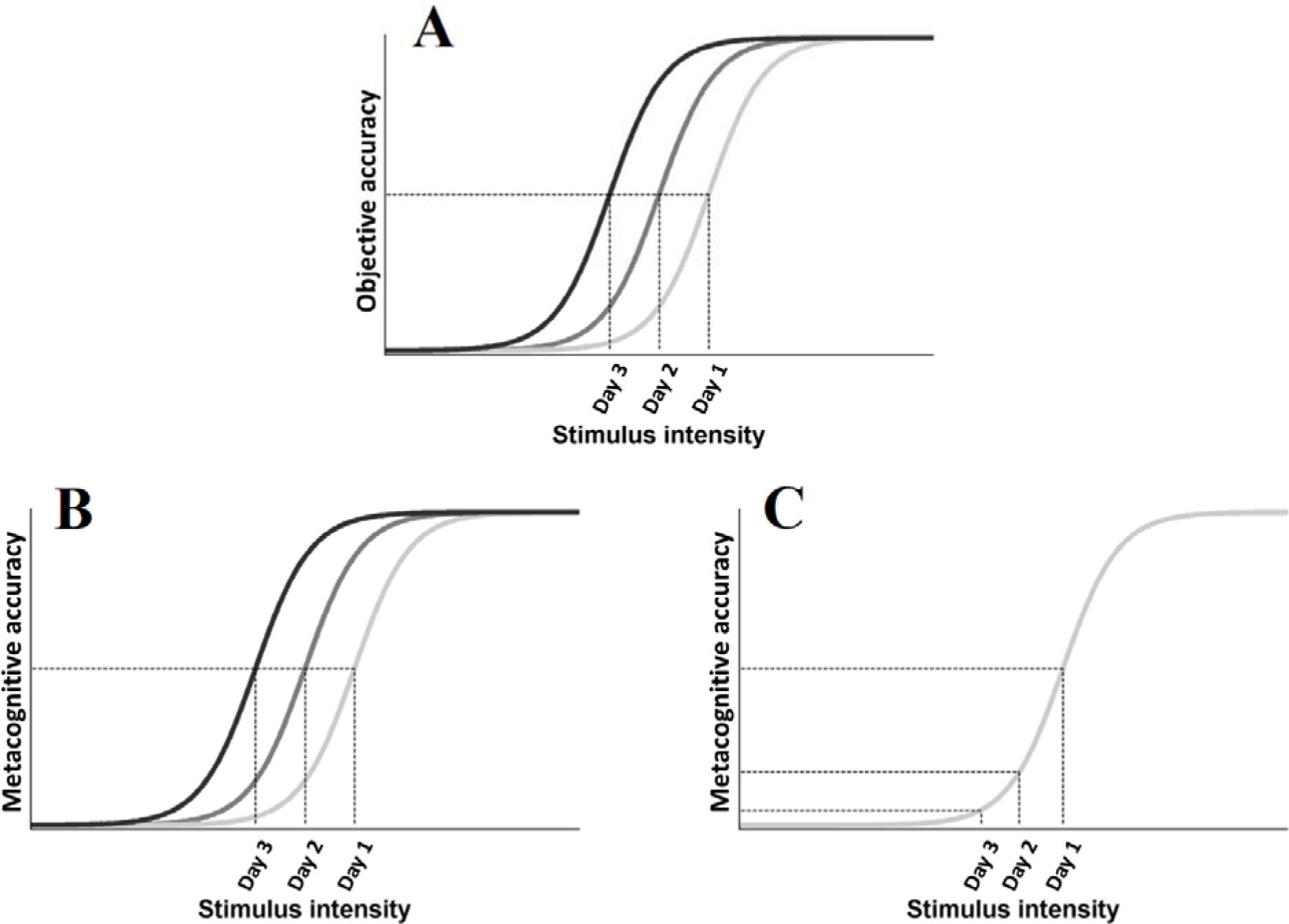
Hypothesized changes in objective and metacognitive psychometric functions with visual perceptual learning (VPL; **a-c**). Psychometric function of objective task accuracy as a function of stimulus intensity (**a**). Psychometric functions of metacognitive accuracy as a function of stimulus intensity (**b & c**), with (**b & c**) corresponding to the presence and absence of improvements in metacognitive accuracy with VPL, respectively. Darker greyscale functions in (**a & b**) correspond to improvements in perception due to VPL across Day 1, 2, and 3. Dashed vertical lines in (**a, b & c**) indicate decreasing stimulus thresholds with VPL across Day 1, 2, and 3, and corresponding objective (**a**) or metacognitive accuracy (**b & c**).

## Experiment 1: Can face identity VPL improve metacognitive accuracy?

### Method

#### *Subject*s

Twenty subjects (13 female & 7 male, *M_age_* = 24.5, *SD* = 5.47) were recruited from Monash University, and reimbursed $60 for their participation. Based on a power analysis (repeated-measures f-test; Faul, Erdfelder, Lang, & Buchner, 2007), sample size was determined to achieve a strong main effect of training on stimulus threshold from day 1 to day 3 (Cohen’s *f* = 0.4, *a* = 0.05, power = 0.95). Subjects reported no history of major medical or psychiatric conditions, and normal or corrected-to-normal vision. All procedures were approved by the Monash University Human Research Ethics Committee (MUHREC), and signed informed consent was obtained prior to the experiment.

#### Stimuli

Four pairs of emotionally neutral, front-facing Caucasian faces (two female & two male) were generated using Facegen software (Singular Inversions, 2006). All faces were converted to greyscale with a black oval mask applied to remove external features (e.g., ears), before normalizing each face pair on their luminance and contrast (Willenbockel et al., 2010). Morpheus software (Morpheus Software, 2014) was then used to morph both faces within a pair together by anchoring key features (e.g., eyes, nose), generating a morph continuum from 100% of one face (100%:0%), to 100% of the other face (0%:100%), in 2% increments. Within the oval mask, faces subtended 9.76 degrees of visual angle (dva) vertically, and 5.9 dva horizontally.

#### Procedure

Subjects performed an unspeeded ABX task across 3 consecutive days. On each trial, subjects judged whether a third face (X) matched the first (A) or second (B) face’s identity, while simultaneously rating the confidence of their decision from 1 (not sure) to 4 (sure) via mouse click (Figure 2A). X was always identical to either A or B, with equal probability. Subjects were encouraged to respond as accurately as possible, and to use the entire confidence scale. No feedback was provided.

All 3 faces (A, B & X) were presented sequentially for 200ms each. To avoid biasing fixations towards a particular facial region (e.g., eyes), each face had a random horizontal leftward or rightward displacement between 0.78 and 1.56 dva, relative to the screen’s centre. The displacement of the first face (A) was randomly determined, with the displacement of the remaining 2 faces (B & X) being opposite to the preceding face, such that only 2 sequences were possible (left (A), right (B), left (X); right (A), left (B), right (X)). Before the presentation of each face, a central leftward- or rightward-pointing arrow (200ms) reliably cued subjects to each faces’ subsequent displacement. After the presentation of each face, a Gaussian noise mask (200ms) appeared which covered the spatial extent of the preceding face.

The task was programmed and run using the Psychophysics toolbox extension for Matlab (Psychtoolbox-3; Brainard, 1997; Pelli, 1997). Stimuli were presented against a grey background on a 23-inch screen (1920x1080 pixels, 60 HZ refresh rate), which subjects viewed from a chinrest placed 75cm away. Subjects were given the chance to take a short break after every 160 trials.

We estimated morph distance thresholds (%) corresponding to 75% accuracy (psychometric slope (β) = 0.1, lapse rate (δ) = 0.05, probability of a correct guessing response (γ) = 0.5), using quick estimation of threshold (QUEST; Watson & Pelli, 1983). QUEST implements an adaptive staircase procedure using Bayesian principles and provides the most probable estimation of stimulus threshold via a posterior distribution function (PDF).

In each trial, threshold estimates (rounded to the nearest multiple of 2%, with a maximum of 50%) were subtracted from, and added to, the morph midpoint (50%:50%) to select A and B faces along the morph continuum. Both faces had an equal probability of corresponding to midpoint±threshold.

The experiment consisted of three phases: Pre-training, training, and post-training (Figure 2C). During both pre- and post-training phases, which occurred immediately before and after training, respectively, subjects performed 2 separate blocks on the ABX task involving all 4 face pairs. Each block consisted of 4 randomly interleaved QUEST staircases. For each face pair, a single staircase (40 trials) was used to independently estimate the morph distance threshold that likely resulted in a discrimination accuracy of 75% within the tested block (i.e., a total of 160 trials/block).

**Fig. 2.**
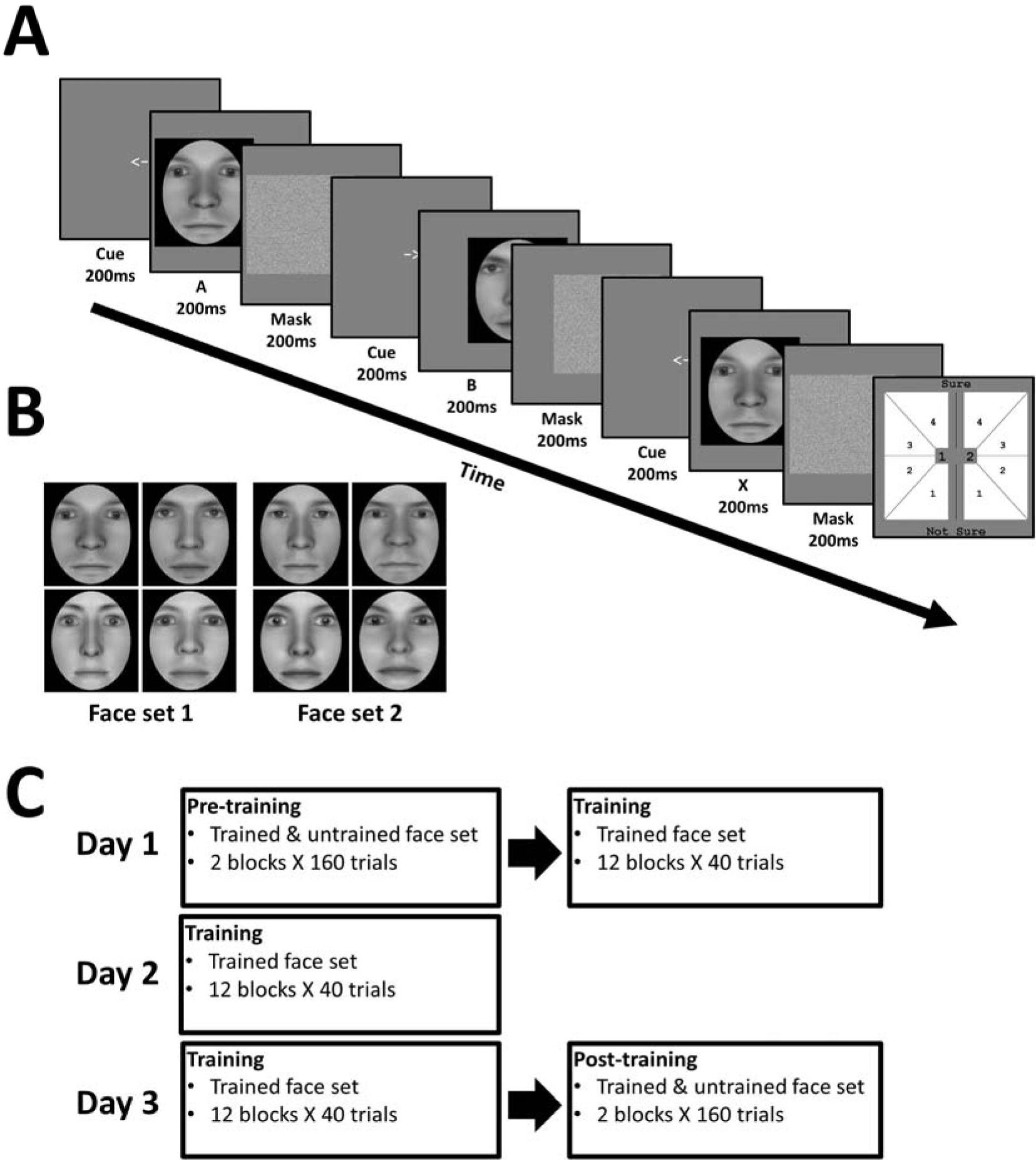
Experiment 1 task (**a**). In each trial, subjects viewed a sequence of three faces (A, B & X). Each face was preceded by a cue corresponding to their on-screen displacement, and followed by a mask. Subjects judged whether the third face (X) matched the identity of the first (A) or second (B) face, corresponding to 1 or 2 in the response screen (last panel), respectively. Subjects also simultaneously reported the confidence of their decision, from 1 (not sure) to 4 (sure). Experiment 1 face sets (**b**). Subjects were trained on one face set, and untrained on the other, in a fully counterbalanced manner. The paradigm for Experiment 1 (**c**) consisted of pre-training, training, and post-training. Training took place across 3 consecutive days, with subjects being trained on a single face set. Pre- and post-training took place immediately before (Day 1) and after (Day 3) training, respectively, with subjects being tested on the trained and untrained face set.

Training occurred across 3 consecutive days, with half of the subjects being trained on face set 1, and the remaining half being trained on face set 2 (Figure 2B). During daily training, subjects performed 12 blocks on the ABX task. Each block consisted of 20 trials for one face pair and 20 trials for the other pair (i.e., a total of 40 trials/block), presented in a randomized order. Within training blocks, the morph distances for both pairs was held constant based on their estimated morph distance threshold. To achieve this, we constructed individual QUEST staircases for each pair based on their previous 80 trials, and the most informative PDF quantile (as determined by QUEST) used to estimate their threshold. To provide threshold estimates for the initial training blocks, subjects performed 2 separate baseline blocks at the beginning of each daily session. On day 1, these corresponded to the pre-training blocks for the trained face pairs. On days 2 and 3, each baseline block instead consisted of 2 interleaved QUEST staircases, with a single staircase (40 trials) for each trained face pair (not shown in Figure 2C).

#### Data analysis

##### Pre- & post-training

Pre- and post-training consisted of 2 blocks. Each block involved 4 randomly interleaved staircases, with a single staircase (40 trials) for each face pair. To measure training effects, we assessed the baseline, pre-training morph distance thresholds for all 4 face pairs. After 3 days of training, we re-assessed the post-training thresholds for all 4 pairs, with subjects being trained on only 2 of these pairs. To achieve this, we constructed individual QUEST staircases (β = 0.1, δ = 0.05, γ = 0.5) based on each face pair’s corresponding 40 trials within pre- and post-training blocks, and defining the mode of the PDF as the threshold (Watson & Pelli, 1983). We then averaged corresponding face pair thresholds to obtain threshold values for the trained and untrained face set for each subject. To test whether post-training thresholds were significantly lower than pre-training thresholds for the trained and the untrained set, we used one-tailed Wilcoxon signed-rank tests.

To estimate the degree of VPL transfer for each subject, we defined a transfer index (TI) as [threshold improvement for the untrained set / threshold improvement for the trained set], with TI = 1 corresponding to complete VPL transfer, and TI = 0 to no transfer (Bi, Chen, Weng, He, & Fang, 2010). We defined threshold improvement as [(Thresholdpre-training – Thresholdpost-training)/Threshold_pre-training_] × 100%. To test whether TI was significantly greater than or less than T1 = 0 and TI = 1, respectively, we used one-tailed one-sample t-tests.

##### Training

Daily training consisted of 12 blocks. Each block involved 20 trials for both trained face pairs presented in a randomized order. Within a given block, the morph distance for either face pair was held constant. For each of the 12 training blocks, we separately calculated threshold, objective accuracy, and metacognitive accuracy, for each pair. For threshold, we constructed individual QUEST staircases (β = 0.1, δ = 0.05, γ = 0.5) based on each pair’s corresponding 20 block trials, and defining the mode of the PDF as the threshold.

We calculated objective accuracy and metacognitive accuracy using a receiver operating characteristics (ROC) curve based on signal detection theory (SDT). For objective accuracy, we constructed a first-order ROC curve relating to the degree of confidence about the perceptual discrimination between face A and B (called Type 1 discrimination in SDT; Macmillan & Creelman, 2005). To achieve this, we considered X=A trials as signal present trials, and X=B trials as signal absent trials. Hits and false alarms were then estimated by systematically varying the SDT criterion in 7 steps. Firstly, we regarded a response as a ‘hit’ when the signal was present, and subjects reported X=A with the highest confidence (4). Similarly, we regarded a response as a ‘false alarm’ when the signal was absent, and subjects reported X=A with the highest confidence (4). We then shifted the criterion to include X=A responses endorsed with a confidence of 3 and 4, and likewise classified responses as hits or false alarms depending on whether the signal was present or absent, respectively. The criterion was shifted in this manner until we obtained hits and false alarms from the highest (4) to lowest (1) confidence ratings for X=A responses, and the lowest (1) to second highest (3) confidence ratings for X=B responses. Plotting hits and false alarms across all 7 criteria produced a ROC curve with 7 inflection points, with the area underneath it, which we term type-I AUC, providing a non-parametric estimate of objective task accuracy (Kaunitz, Rowe, & Tsuchiya, 2016; Macmillan & Creelman, 2005; Wilimzig, Tsuchiya, Fahle, Einhauser, & Koch, 2008).

For metacognitive accuracy, we constructed a second-order ROC curve relating to the degree of confidence about correct and incorrect perceptual decisions. To achieve this, we considered trials where perceptual decisions were correct (i.e., subjects reported X=A in X=A trials, and X=B in X=B trials) as signal present trials, and trials where perceptual decisions were incorrect (i.e., subjects reported X=B in X=A trials, and X=A in X=B trials) as signal absent trials. Hits and false alarms were then estimated by systematically varying the SDT criterion in 3 steps. Firstly, we regarded a response as a ‘hit’ when a signal present trial was endorsed with the highest confidence (4), and a ‘false alarm’ when a signal absent trial was endorsed with the highest confidence (4). We then shifted the criterion to include responses endorsed with a confidence of 3 and 4, and likewise classified responses as hits or false alarms depending on whether the signal was present or absent, respectively. The criterion was shifted in this manner until we obtained hits and false alarms from the highest (4) to second lowest (2) confidence ratings. Plotting hits and false alarms across all 3 criteria produced a ROC curve with 3 inflection points, with the area underneath it, which we term type-II AUC, providing a non-parametric estimate of metacognitive accuracy (Galvin et al., 2003; Kaunitz et al., 2016; Wilimzig et al., 2008).

To investigate main effects of daily training on morph distance threshold, type-I AUC, and type-II AUC, we performed separate linear mixed-effects analyses using lme4 package (Bates, Mächler, Bolker, & Walker, 2015) within R (R Development Core Team, 2014). We constructed a 2x3 nested mixed design, with trained face set (i.e., face set 1 or face set 2) as a between-subject variable, and daily training session (i.e., 1st to 3rd day) as a within-subject variable. We modelled daily training session as a fixed effect. For our random effects, we modelled an intercept for face set to account for variances in learning effects between both sets. Furthermore, a by-subject intercept and slope was modelled for daily training session to account for subject variability in learning effects and the rate of these effects across training sessions. We performed likelihood ratio tests between the full model, as described above, and a reduced model, without daily training sessions modelled as a fixed effect, to obtain chi-squared statistics and associated p-values.

### Results and Discussion

The results of Experiment 1 are shown in Figure 3. From pre- to post-training, morph threshold significantly decreased for both the trained (*Z* = −3.51, *p* < 0.001) and the untrained (*Z* = −3.32, *p* < 0.001) face set (Figure 3A). Quantifying this transfer, transfer index (TI, see method) was 0.67 (*SD* = 1.20), which was significantly greater than 0 (*p* = .011), but not less than 1 (*p* = .11). This suggests that with training, subjects successfully demonstrated higher-order, identity-invariant VPL of face identity.

**Fig. 3.**
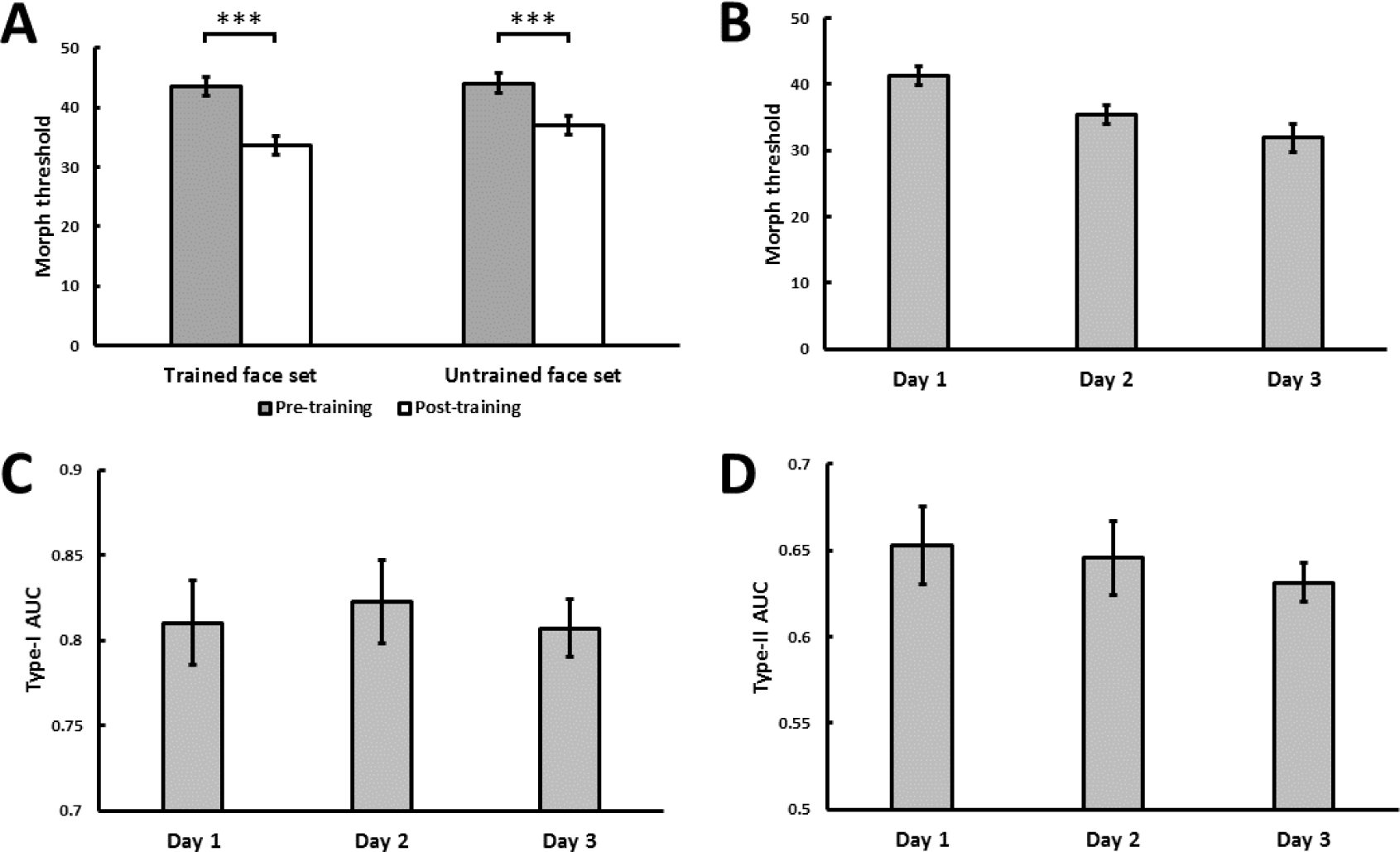
Experiment 1 face identity VPL (**a-d**). Training effects measured as median morph threshold for trained and untrained face sets before (on Day 1) and after (on Day 3) training sessions (**a**). Asterisks indicate significant differences (*** *p* < .001). (**b-d**) Mean training effects measured as threshold (**b**), type-I AUC for objective accuracy (**c**), and type-II AUC for metacognitive accuracy (**d**), as a function of daily training. All 3 variables were independently estimated within each training block (see Method). Error bars denote ± 1 within-subjects SEM (Cousineau, 2005).

With daily training, morph threshold steadily decreased, confirmed by a significant main effect of training on threshold (*χ*^2^(1) = 27.06, *p* < .001; Figure 3B). As intended by our QUEST procedure however, no main effect of training on objective accuracy, measured as type-I AUC, was observed (*χ*^2^(1) = 0.10, *p* > .250; Figure 3C). Importantly, metacognitive accuracy, measured as type-II AUC, decreased with training (*χ*^2^(1) = 4.52, *p* = .034; Figure 3D). This result is consistent with our second hypothesis (Figure 1C), suggesting that although subjects learned to discriminate faces based on their identity, they were unable to consciously access the source of this learning. If they were able, their metacognitive accuracy should have tracked their objective accuracy (Figure 1B), which was held constant by our QUEST procedure (Figure 3C). Together, our findings support a model such as reverse hierarchy theory (Ahissar & Hochstein, 2004; Hochstein & Ahissar, 2002) where, in the absence of top-down guided VPL (e.g., VPL of high-level stimulus properties), the source of VPL (e.g., spatial relationship between stimulus elements) remains consciously inaccessible. However, this result can also be accounted for by a VPL model which argues that, regardless of task-relevant stimulus complexity, learned stimuli should remain consciously inaccessible (e.g., Squire & Wixted, 2011). Thus, we conducted a second experiment to investigate whether or not the VPL of face contrast, a low-level face property, can improve the metacognitive accuracy of perceptual judgements.

## Experiment 2: Can face contrast VPL improve metacognitive accuracy?

### Method

#### *Subject*s

Twenty subjects (14 female & 6 male, *M_age_* = 24.9, *SD* = 5.24), who did not participate in Experiment 1, were recruited from Monash University, and reimbursed $60 for their participation. Subjects reported no history of major medical or psychiatric conditions, and normal or corrected-to-normal vision. All procedures were approved by MUHREC, and signed informed consent was obtained prior to the experiment.

#### Stimuli and Procedure

Four emotionally neutral, front-facing Caucasian faces (2 female & 2 male) that differed in identity to the faces used in Experiment 1, were generated using FaceGen software. Using the same method as Experiment 1, each face was converted to greyscale, and a black oval mask applied. Experiment 2 followed the same procedure as Experiment 1, with the exception of the following changes. In each trial of the ABX task, the same face was presented as faces A, B, and X (i.e., identity remaining constant in a given trial). The contrast of A, B, and X, defined as their normalized root mean square (nRMS) contrast, was manipulated. Subjects were instructed to judge whether the third face (X) matched the contrast of the first (A) or second (B) face (Figure 4A). The contrast of X was always identical to either A or B, with equal probability.

**Fig. 4.**
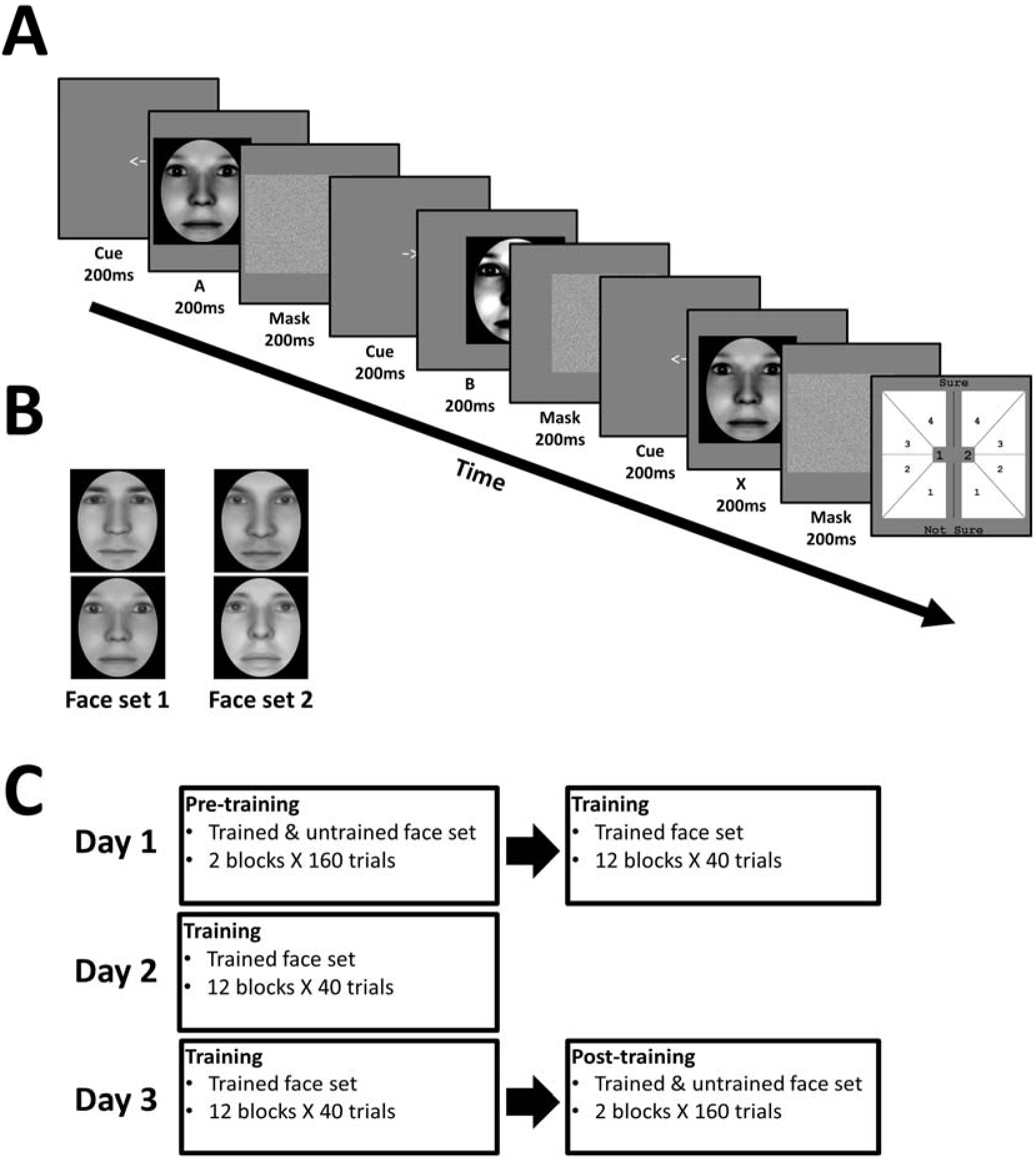
Experiment 2 task, which is identical to Experiment 1, except for the nature of the face stimuli (**a**). Experiment 2 face sets (**b**). Subjects were trained on one face set, and untrained on the other, in a fully counterbalanced manner. The paradigm for Experiment 2 (**c**) was identical to the paradigm of Experiment 1.

To obtain nRMS contrast, we calculated the standard deviation of contrast within the oval mask of each face, and normalized it by their mean luminance (set to 125 cd/m^2^ for all faces). We chose nRMS contrast based on its reliability in predicting human contrast sensitivity to natural images (Bex & Makous, 2002).

We used QUEST to estimate nRMS contrast different thresholds in log scale corresponding to 75% accuracy (β = 3.5, δ = 0.05 & γ = 0.5). In each trial, we first converted threshold estimates to linear scale (i.e., 10^Threshold^), and then subtracted it from, and added it to, the nRMS contrast midpoint (0.5 linear scale) to derive the contrast values for A and B faces. Both faces had an equal probability of corresponding to midpoint±threshold.

Both pre- and post-training phases involved all 4 faces. During training, half of the subjects were trained on face set 1, and the remaining half were trained on face set 2 (Figure 4B; Figure 4C).

#### Data analysis

Experiment 2 followed the same data analysis as Experiment 1, with the following changes. For pre-training, training, and post-training, constructed QUEST staircases used the following parameters: β = 3.5, δ = 0.05 and γ = 0.5. Furthermore, contrast threshold estimates from each staircase were converted to linear scale (i.e., 10^Threshold^) prior to formal statistical analyses.

### Results and Discussion

The results of Experiment 2 are shown in Figure 5. From pre- to post-training, contrast threshold significantly decreased for both the trained (*Z* = −2.17, *p* = 0.015) and untrained (*Z* = −2.54, *p* = 0.006) face set (Figure 5A). Quantifying this transfer, transfer index (TI, see Exp. 1 method) was 0.62 (*SD* = 1.25), which was significantly greater than 0 ( *p* = .02), but not less than 1 (*p* = 0.09). This suggests that with training, subjects successfully demonstrated higher-order, identity-invariant VPL of face contrast.

**Fig. 5.**
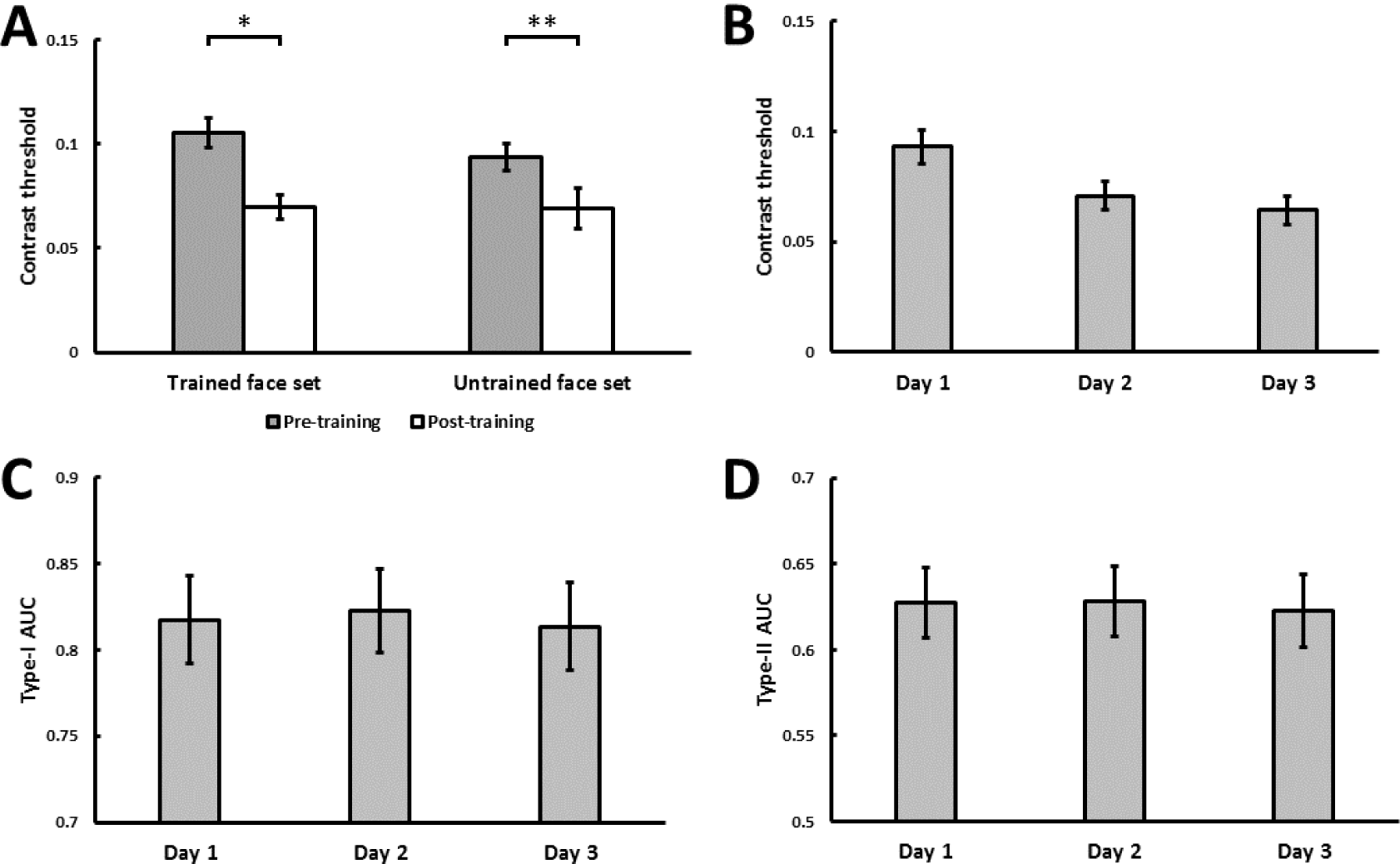
Experiment 2 face contrast VPL (**a-d**). Training effects measured as median contrast threshold for trained and untrained face sets before (on Day 1) and after (on Day 3) training sessions (**a**). Asterisks indicate significant differences (* *p* < .05, \*\**p* < .01). (**b-d**) Mean training effects measured as threshold (**b**), type-I AUC for objective accuracy (**c**), and type-II AUC for metacognitive accuracy (**d**), as a function of daily training. All 3 variables were independently estimated within each training block (see Method). Error bars denote ± 1 within-subjects SEM (Cousineau, 2005).

With daily training, contrast threshold steadily decreased, confirmed by a significant main effect of training on threshold (*χ*^2^(1) = 12.62, *p* < .001; Figure 5B). As intended by our QUEST procedure however, no main effect of training on objective accuracy, measured as type-I AUC, was observed (*χ*^2^(1) = .28, *p* > .250; Figure 5C). Importantly, no main effect of training on metacognitive accuracy, measured as type-II AUC, was also found (*χ*^2^(1) = .34, *p* > .250; Figure 5D). This result is consistent with our first hypothesis (Figure 1B), suggesting that as subjects learned to discriminate faces based on their contrast, they were able to consciously access the source of this learning. If they were unable, metacognitive accuracy should have dissociated from objective accuracy (Figure 1C), which was held constant by our QUEST procedure. Thus, our finding suggests that the VPL of face contrast accompanies improved conscious accessibility to the source of VPL.

## General Discussion

We sought to investigate whether or not the metacognitive accuracy of perceptual judgements could be improved by VPL. Holding objective accuracy constant, metacognitive accuracy was not improved by face identity VPL (Figure 3), but was improved by face contrast VPL (Figure 5). Together, this suggests that improved conscious stimulus accessibility is absent with the VPL of a high-level face property, and present with the VPL of a low-level face property. To our knowledge, this is the first study to obtain such a counterintuitive and dissociative finding, given that conscious stimulus processing has been associated with the VPL of high- (Axelrod & Rees, 2014) but not low-level (Seitz et al., 2009; Watanabe et al., 2001) stimulus properties.

Our finding is inconsistent with the view that VPL should not improve conscious accessibility (e.g., Squire & Wixted, 2011). At least one form of VPL, namely face contrast VPL, cannot be explained by such models. Rather, our main findings can be interpreted in a model similar to reverse hierarchy theory (RHT; Ahissar & Hochstein, 2004; Hochstein & Ahissar, 2002). RHT however, has not made explicit predictions about whether metacognitive accuracy should or should not be improved by the VPL of high- or low-level properties. Thus, we propose our following ‘RHT-like’ model to be distinguished from original RHT (Ahissar & Hochstein, 2004; Hochstein & Ahissar, 2002).

According to our RHT-like model, conscious accessibility operates in reverse of the visual hierarchy alongside VPL. Specifically, VPL proceeds by modifying high-level stimulus representations via bottom-up processing, and if insufficient (e.g., poor spatial resolution), regresses down the hierarchy and modifies lower-level representation via top-down processing. Similarly, conscious accessibility begins at a stimulus’ gist via bottom-up processing, and regressing down the hierarchy to its lower-level elements via top-down processing. Thus, in the absence of top-down guided VPL (e.g., VPL of high-level properties), the source of VPL cannot be consciously accessed. This manifested as the dissociation between improved objective accuracy and non-improved metacognitive accuracy with face identity VPL (Figure 3C & 3D), which is consistent with non-improved conscious accessibility (Figure 1C). While our study cannot explain what source of VPL remains consciously inaccessible, one possibility is the precise spatial relationship between facial elements, although future research is needed to clarify this. In contrast, in the presence of top-down guided VPL (e.g., VPL of low-level properties), the source of VPL can be consciously accessed. This was demonstrated in the concordant improvement in objective accuracy and metacognitive accuracy with face contrast VPL (Figure 5C & 5D), which is consistent with improved conscious accessibility (Figure 1B). This finding implies that greater contrast sensitivity is accompanied with increasing conscious accessibility. Future studies however, are needed to localise the neural networks supporting such improvements in conscious accessibility.

In addition to providing insights into how VPL shapes conscious accessibility, our study can inform future research by addressing two limitations within the literature. Firstly, VPL studies of metacognition have largely failed to control objective accuracy (e.g., Bertels et al., 2012; Schlagbauer et al., 2012; Schwiedrzik et al., 2011), which typically correlates with measures of metacognition (Galvin et al., 2003; Sandberg et al., 2010). Without using our staircase procedure to control this confound, we may have failed to dissociate objective accuracy from metacognitive accuracy in Experiment 1. Secondly, in clinical VPL studies, especially within amnesic populations (e.g., Fahle & Daum, 2002; Manns & Squire, 2001), the absence of conscious accessibility has been inferred from intact VPL. Measuring metacognitive accuracy however, we found evidence suggesting that the presence of VPL does not necessarily indicate the absence of conscious accessibility. Thus, extending our approach to both healthy and psychiatric populations can provide further insights into how VPL shapes conscious accessibility.

In conclusion, we found evidence indicating that conscious accessibility is improved by the VPL of a low- but not high-level face property. Outside VPL, our study can open new avenues to explore the relationship between metacognition and other learning paradigms such as classical (Pearson, 2012) and operant (Pessiglione et al., 2008) conditioning, as well as learning concepts such as artificial grammar (Scott, Dienes, Barrett, Bor, & Seth, 2014). Understanding the relationship between learning and consciousness in turn, constrains how our conscious experience is shaped by learned information, a central question in cognitive neuroscience (Bayne et al., 2009).

## Author Contributions

BC and NT developed the study concept and design. BC performed testing and data collection. BC performed the data analysis and interpreted the results under the supervision of NT. BC and NT drafted the manuscript. MM provided comments on the manuscript. All authors approved the final version for submission.

## Acknowledgements

We are very grateful to Lisandro Kaunitz and Roger Koenig for their Matlab programming advice. N.T. was supported by Precursory Research for Embryonic Science and Technology project from the Japan Science and Technology Agency (3630), the Future Fellowship (FT120100619) and the Discovery Project (DP130100194) from the Australian Research Council.

